# A Triple-Modality Peptide-Antibiotic-Phage Therapy Eradicates Multidrug-Resistant *Serratia marcescens* Biofilms

**DOI:** 10.64898/2026.04.08.717253

**Authors:** Aryaan P. Duggal, Adit B. Alreja, Isha Vashee, Hayley Nordstrom, Erin Harrelson, Nakia Fallen, Kari-Ann Takano, Ryan A. Blaustein, Derrick E. Fouts, Norberto Gonzalez-Juarbe

**Affiliations:** University of Maryland, College Park, Department of Cell Biology and Molecular Genetics; Human Genomic Medicine, J. Craig Venter Institute.; University of Maryland, College Park, Department of Nutrition and Food Science.

## Abstract

*Serratia marcescens* is an opportunistic pathogen that causes severe hospital-acquired infections, notable for its biofilm formation abilities and development of extensive antibiotic resistance. Here we evaluated the efficacy of bacteriophages, antibiotics, and antimicrobial peptides (BAP), alone and in combination, against fourteen multi-drug-resistant (MDR) *S. marcescens* isolates sourced from hospitals and other environmental settings in an *in vitro* biofilm model. Phage combination with a cocktail of sub-minimal inhibitory concentration (MIC) of penicillin-streptomycin, kanamycin, and ciprofloxacin, reduced biofilm biomass, however, complete decolonization was not achieved. Incorporating an antimicrobial peptide cocktail into this regimen eradicated 99.99% of multi-drug-resistant isolates grown planktonically or in surface-associated biofilms. Microscopy and viability assays confirmed extensive biofilm disruption and bacterial clearance without regrowth. These findings reveal that simultaneous interference of cell wall synthesis, protein translation, DNA replication, and membrane integrity can overcome *S. marcescens* antimicrobial defenses, establishing a multifaceted therapeutic framework for managing device-associated infections caused by MDR pathogens.

## Introduction

*Serratia marcescens* is a Gram-negative, facultative anaerobic bacterium within the Enterobacterales order. Historically considered an environmental organism, with reservoirs spanning soil, plants, and surface water, it has become a widespread opportunistic pathogen responsible for numerous hospital-associated infections, particularly in immunocompromised patients, the elderly, and neonates^1^. The ability of *S. marcescens* to establish biofilms and persist in hospital environments such as on medical equipment has led to frequent outbreaks in intensive care units (ICUs) and neonatal intensive care units (NICUs) which has proven to be deadly to the occupying patients^2,3^. Epidemiological studies have documented the increasing burden of *S. marcescens* infections in healthcare settings^4,5^. Notably, neonatal and geriatric patients are disproportionately affected by nosocomial infections, including sepsis caused by *S. marcescens*. These patients suffer from higher mortality rates, longer-term morbidity, and increased cost of care^6^. These resulting complications are significantly worsened in strains of *S. marcescens* that exhibit resistance to broad-spectrum antibiotics^7,8^.

*S. marcescens* exhibits both intrinsic and acquired antibiotic resistance mechanisms, complicating treatment strategies. Intrinsically, the bacterium possesses efflux pumps and reduced outer membrane permeability, limiting antibiotic uptake^7^. Acquired resistance arises through horizontal gene transfers and mutations affecting antibiotic targets. For example, mutations in DNA gyrase (*gyrA, gyrB*) can confer resistance to fluoroquinolones, while ribosomal modifications reduce aminoglycoside efficacy^9,10^. A study conducted by Bryant et. al showed *S. marcescens* gained a bla_KPC-4_ gene from *Enterobacter cloacae* via a plasmid through a wound in a hospital acquired infection of a patient, highlighting the acquisition of antimicrobial resistance genes ^11^ In addition, some strains of *S. marcescens* can produce β-lactamases that degrade β-lactam antibiotics, including penicillins and cephalosporins^12^. *S. marcescens* environmental persistence and acquisition of antimicrobial resistance genes is enhanced by a defining virulence trait in its ability to form biofilms, which are structured bacterial communities surrounded by an extracellular matrix composed of polysaccharides, proteins, and extracellular DNA^13^. Biofilm formation facilitates colonization of inanimate surfaces, particularly moist areas such as sinks and drains but also medical devices such as central venous catheters, endotracheal tubes, and ventilators^14,15^. Within biofilms, bacteria exhibit an increased resistance to host immune responses and antimicrobial agents^16,17^. Studies indicate that biofilm-associated *S. marcescens* infections require antibiotic concentrations significantly higher than planktonic infections, often exceeding clinically feasible doses to safely use within patients^8,18^. The resilience of *S. marcescens* biofilms and that of other Enterobacterales members pose significant challenges for hospital sanitation protocols such as with standard disinfectants including bleach, ammonium chloride and other chlorine-based cleaning agents^15^. Moreover, biofilms can persist on surfaces for extended periods, even under stringent cleaning regimens, contributing to recurrent outbreaks in healthcare settings^4,14,19–22^. Taken together, it is imperative to design new strategies to decolonize biofilms and mitigate risks for the spread of *S. marcescens* in the hospital setting.

Despite adherence to standard sanitation protocols, current disinfection strategies can be inadequate in the eradication of *Serratia* spp. biofilms in healthcare environments, necessitating the need for development of novel approaches. Widely used agents such as diluted bleach exhibit limited efficacy against mature biofilms due to poor penetration of the extracellular matrix and rapid neutralization in organic-rich environments^23^. Similarly, quaternary ammonium compounds (QACs) have been shown to reduce planktonic bacterial loads while leaving biofilm-embedded organisms largely viable^24^. Biocide and antibiotic resistance genes frequently occur together and biocide chemical residues on environmental surfaces may select for “qac” antimicrobial resistance genes^25^. Hydrogen peroxide vapor systems, though more effective in some contexts, require prolonged exposure and environmental containment to exert bactericidal effects, which limits their practical utility in high-turnover clinical settings^26^. *In situ* investigations have confirmed the persistence of biofilm-forming *S. marcescens* strains on surfaces that were considered disinfected, contributing to repeated outbreaks in NICUs and intensive care units^27^. Antibiotic lock therapy in combination with systemic antibiotic therapy has been recently found to be an effective way to treat peripherally inserted central venous catheter-related bloodstream infections^28^.

However, complexity of the biofilm’s organic architecture, combined with the bacterium’s innate and acquired resistance mechanisms, renders most standard sanitation efforts insufficient. These limitations underscore the need for engineered combination therapies that are multi-faceted in their mechanism of eradication and employ a multi-drug approach to effectively eliminate *S. marcescens*.

In quest of such multi-faceted therapies, we set out to screen the effectiveness of major antibiotics in their ability to kill multidrug-resistant (MDR) *S. marcescens* both in planktonic and biofilm form. We found different classes of antibiotics exhibited varying levels of killing activity on planktonic and biofilm associated *S. marcescens*. A combination regimen using sub-MIC concentrations of an antibiotic cocktail showed dose-dependent efficacy at reducing bacterial titers in planktonic form and in causing a reduction in biofilm biomass. As natural predators of bacteria, phages have evolved to selectively kill target bacteria by hijacking their biological machinery. A growing body of evidence suggests that in some situations, phages can be utilized synergistically with antibiotics to improve antimicrobial effectiveness. Our results showed that a combination of bacteriophage and antibiotics reduced biofilm biomass to a greater degree than the antibiotic mixture alone, highlighting the synergistic effect of phage therapy and antibiotic treatment. To take our findings further, we employed a Bacteriophage + Antibiotics + Antimicrobial Peptides (BAP) approach and found this triple treatment was the most effective at decolonizing biofilms especially on MDR *S. marcescens*. Taken together, our findings highlight the importance of multidimensional antimicrobial approaches to target *S. marcescens* associated infections.

## Results

### Distinct antibiotic classes exert variable inhibitory effects on *S. marcescens* in planktonic and biofilm forms

First, we wanted to define the antimicrobial resistance properties of *S. marcescens* in planktonic form against existing antibiotics. To define the antimicrobial resistance properties, we used antibiotics that target different bacterial components on the *S. marcescens* UT-383 strain, such as cell wall synthesis (β-lactams), protein synthesis (aminoglycosides, tetracyclines, and chloramphenicol), DNA replication (fluoroquinolones), and folate synthesis (sulfonamides) (**Table 1**). Treatment with β-lactam and aminoglycoside containing drugs Penicillin-Streptomycin and ampicillin at concentrations of 0.80 mg/mL and 0.08 mg/mL resulted in a substantial reduction in bacterial density, with Penicillin-Streptomycin exhibiting a lower OD_600_ at both concentrations and Ampicillin a less pronounced effect (**Fig. 1A**). For aminoglycosides and tetracyclines, Kanamycin and Tetracycline at 0.80 mg/mL caused a large reduction in bacterial titers, while at 0.08 mg/mL the reduction was not significant. Gentamicin was observed to partially decrease the titers of *S. marcescens* at both doses (**Fig. 1B**). As previously observed by our group^8^, Chloramphenicol (an amphenicol) displayed dose-dependent inhibition of *S. marcescens* (**Fig. 1C**). Fluoroquinolone and Antifolate, Ciprofloxacin and Trimethoprim, respectively, also showed a dose-dependent decrease of bacterial numbers with the higher dose providing over 50% reduction (**Fig. 1D, E**). These findings reveal a clear pattern of variable susceptibility among planktonic S. *marcescens* to different antibiotic classes, underscoring the importance of employing combination strategies against bacterial targets.

**Figure 1:**
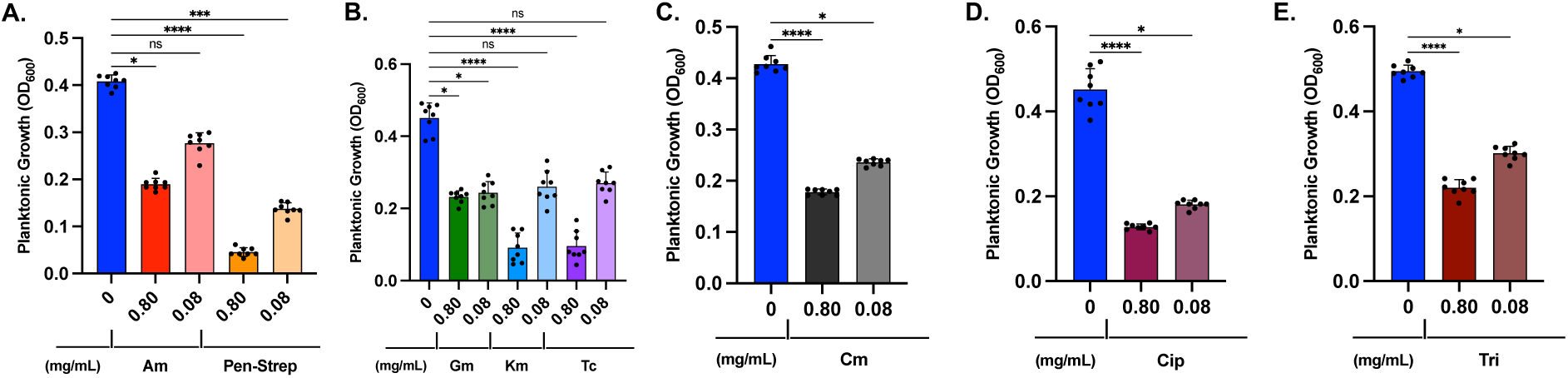
Major antibiotic classes have differential efficacy against planktonic *S. marcescens*. *Serratia marcescens* UT-383 was grown planktonically to mid-Log phase followed by treatment with Ampicillin, Penicillin-Streptomycin, Gentamicin, Kanamycin, Tetracycline, Chloramphenicol, Ciprofloxacin, and Trimethoprim for 24 hours followed by absorbance reading at OD600. Statistical differences were determined by the Kruskal–Wallis test with Dunn’s multiple-comparison post-test. * p ≤ 0.05, ** p ≤ 0.01, *** p ≤ 0.001, **** p ≤ 0.0001. Error bars denote standard deviation.

Biofilms are known for providing bacteria with formidable resilience against antimicrobial agents and relevance to device-associated infections posing a greater therapeutic challenge^14,15,17^. Given this knowledge, we aimed to evaluate the activity of the different classes of antibiotics against highly structured mature biofilms. Using the high (0.80 mg/mL) and low (0.08 mg/mL) doses of antibiotics as in **Fig. 1**, on *S. marcescens* UT-383, we observed that Penicillin-Streptomycin and Ciprofloxacin (**Fig. 2A, D**) were the most effective at each dose in reducing biofilm biomass. Ampicillin, Chloramphenicol, Kanamycin and Tetracycline (**Fig. 2A-C**) showed a dose dependent effect, and Gentamicin along with Trimethoprim only partially reduced biofilm biomass (**Fig. 2B, E**).

**Figure 2:**
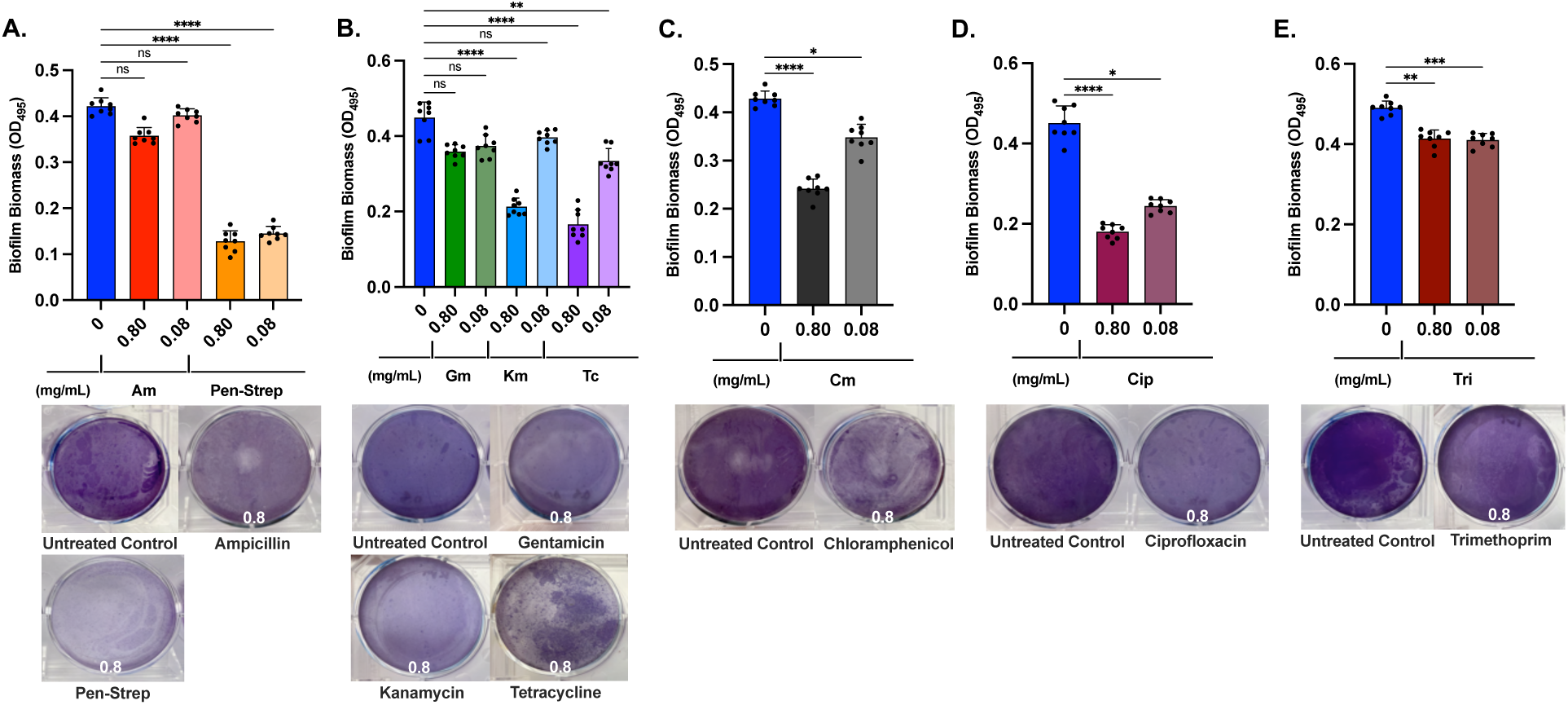
Major antibiotic classes have differential efficacy against mature *S. marcescens* biofilms. *Serratia marcescens* UT-383 was grown in biofilm form for 48 hours prior to treatment with Ampicillin, Penicillin-Streptomycin, Gentamicin, Kanamycin, Tetracycline, Chloramphenicol, Ciprofloxacin, and Trimethoprim. Statistical differences were determined by the Kruskal–Wallis test with Dunn’s multiple-comparison post-test. * p ≤ 0.05, ** p ≤ 0.01, *** p ≤ 0.001, **** p ≤ 0.0001. Error bars denote standard deviation.

### Combination of sub-MIC concentrations of antibiotics are effective against *S. marcescens* in planktonic and biofilm forms

Given the variation in antibiotic susceptibility observed, we sought to employ a sub-MIC antibiotic combination regimen over a single high-dose treatment. Since antibiotics also destroy commensal bacteria, a multi-drug, lower dose approach could potentially lessen these harmful effects on commensals. We combined Penicillin-Streptomycin, Ciprofloxacin, and Kanamycin to target multiple bacterial pathways at sub-MIC concentrations. Penicillin-Streptomycin disrupts cell wall and protein synthesis, Ciprofloxacin inhibits DNA gyrase and topoisomerase IV, and Kanamycin halts translation via inhibition of the 30S ribosomal subunit^29^ (**Fig. 3A**). This multi-pronged strategy was validated by challenging the planktonic bacteria in the Log phase with diverse doses of the antibiotic cocktail comprised of thirds for each Penicillin-Streptomycin, Ciprofloxacin, and Kanamycin. The highest dose of 100 µg/mL (33.3 µg/mL of each antibiotic) antibiotic mixture reduced bacterial titers to near background levels, while 10-fold dilutions to 10 µg/mL, 1 µg/mL, 0.1 µg/mL and 0.01 µg/mL showed a dose dependent reduction in efficacy with only 10 µg/mL and 1 µg/mL significantly reducing planktonic bacterial titers (**Fig. 3B**).

**Figure 3:**
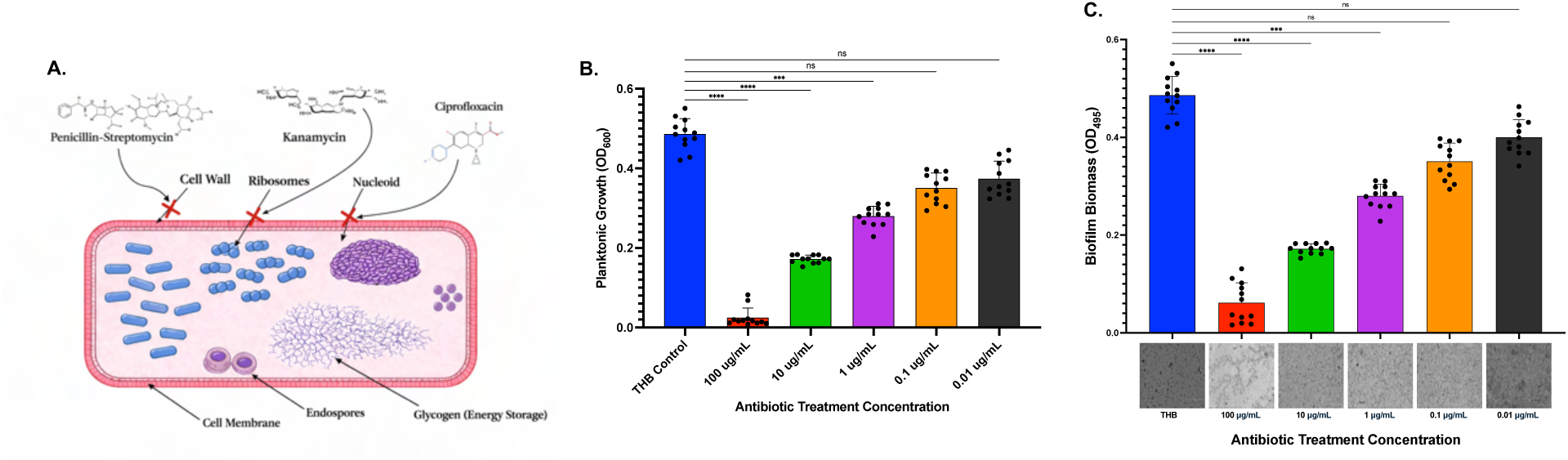
Sub-MIC antibiotic cocktail leads to decreased planktonic growth and biofilm biomass in a dose dependent manner. **A**. Schematic of antimicrobial targets of selected antibiotics. **B**. Planktonic grown bacteria were challenged mid-log with a combination mixture composed of three antibiotics (PenStrep, ciprofloxacin, and kanamycin) at concentrations between 100 µg/mL to 0.01 µg/mL and absorbance at OD600 was measured after 24 hours. **C**. Biofilm biomass measured from crystal violet staining of mature biofilms treated with the antibiotic at concentrations between 100 µg/mL to 0.01 µg/mL for 24 hours prior to staining and OD 495 absorbance reading. Statistical differences were determined by the Kruskal–Wallis test with Dunn’s multiple-comparison post-test. * p ≤ 0.05, ** p ≤ 0.01, *** p ≤ 0.001, **** p ≤ 0.0001. Error bars denote standard deviation.

To test the effectiveness of our antibiotic mixture against *S. marcescens* biofilms, we generated static biofilms and challenged them with 5 different doses of the antibiotic mixture ranging from 0.01 to 100 µg/mL as was done for planktonic bacteria. Here we observed that 100 and 10 µg/mL of our antibiotic mixture resulted in a major reduction in biofilm biomass. The 1 µg/mL, 0.1 µg/mL and 0.01 µg/mL doses had a dose dependent reduction in effectiveness against *S. marcescens* biofilm. Microscopy images mirrored these results, showing disrupted biofilm architecture at higher concentrations and intact structures at lower doses (**Fig. 3C**). Taken together, this data suggests that a sub-MIC antibiotic mixture is sufficient to disrupt planktonic and biofilm bacterial cultures of *S. marcescens*.

### Combination of *Serratia marcescens* bacteriophage and antibiotic cocktail leads to enhanced biofilm disruption

While sub-MIC antibiotic strategies have been explored to mitigate resistance development, their efficacy remains limited due to incomplete bacterial clearance and the potential for promoting adaptive resistance mechanisms. Consequently, alternative antimicrobial approaches are needed, with bacteriophages emerging as a promising strategy. We identified two bacteriophages (ϕJCVI_Sm1 and ϕJCVI_Sm2) from municipal wastewater that produced plaques on our laboratory MDR *S. marcescens* strain UT-383 and a non-MDR strain HER1170, grown planktonically on agar plates (**Supplemental Fig. 1A-B**). To determine the efficacy of these phages on mature *S. marcescens* biofilms, we tested each phage at MOIs of 0.01, 0.1, 1 and 10. We observed that ϕJCVI_Sm2-challenged biofilms was slightly more efficient in reducing biofilm biomass than ϕJCVI_Sm1 at lower MOIs (**Supplemental Fig. 1C**). When we tested ϕJCVI_Sm2 alone and in combination with our antibiotic mixture, the results revealed that the combination of MOI 0.1 phage with 10 µg/mL antibiotic mixture reduced biofilm biomass by 88%, compared to a 63% reduction achieved by the antibiotic mixture alone (**Fig. 4**). Microscopy confirmed that biofilm architecture was severely disrupted in combined treatments, while completely intact biofilm architecture was only present within the single treatments. These results highlight the synergistic effects of phage therapy and antibiotic combinations on biofilm disruption.

**Figure 4:**
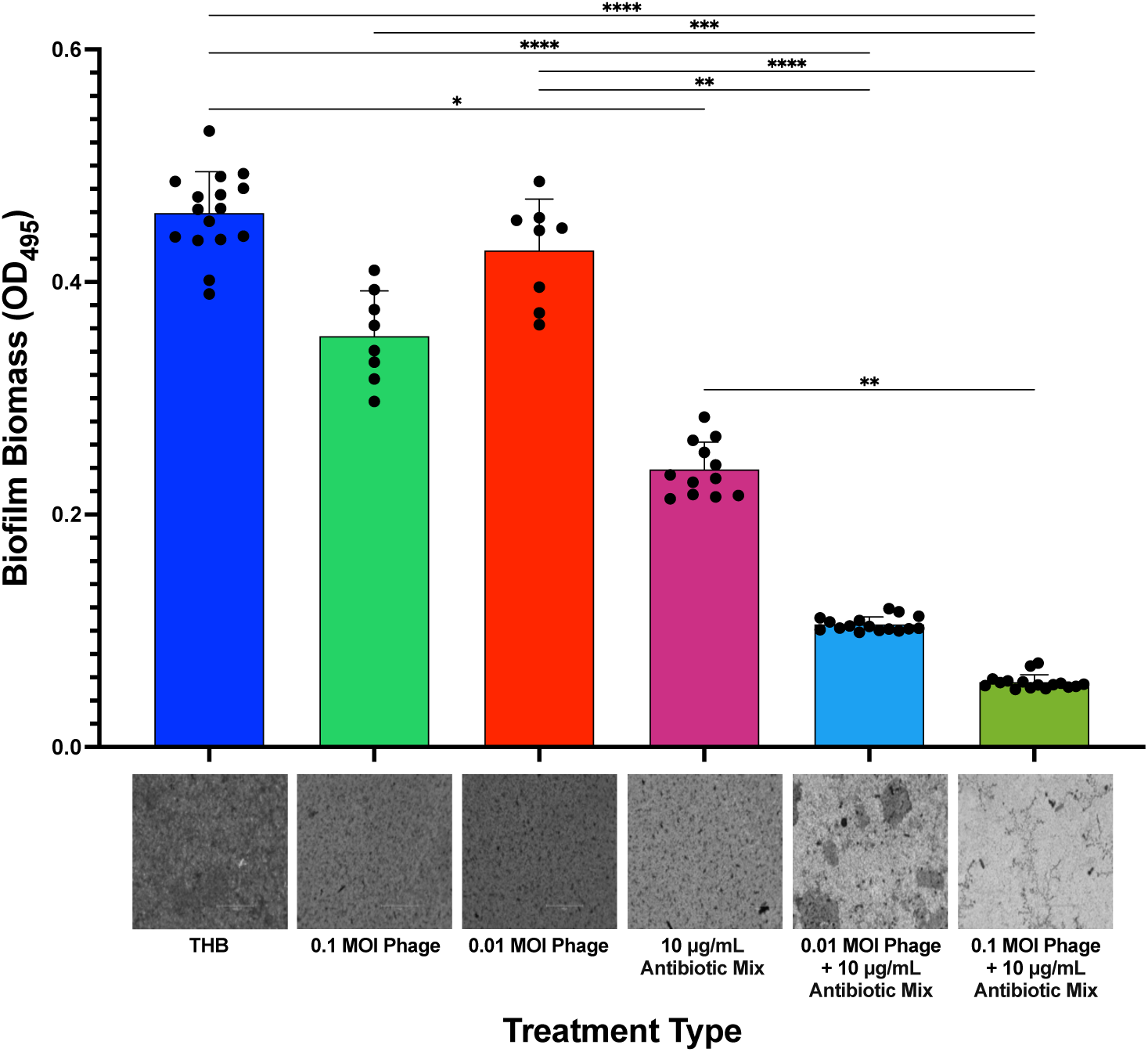
Antibiotic-bacteriophage combination has a synergistic effect against *S. marcescens* biofilms. Biofilm biomass quantification following *S. marcescens* UT-383 treatment the with the antibiotic cocktail totaling 10 µg/mL, phage at MOI 0.1 or 0.01 or combinations at the same doses. Crystal violet staining was used to assess biofilm biomass post treatment. Statistical differences were determined by the Kruskal–Wallis test with Dunn’s multiple-comparison post-test. * p ≤ 0.05, ** p ≤ 0.01, *** p ≤ 0.001, **** p ≤ 0.0001. Error bars denote standard deviation.

### MDR *S. marcescens* isolates have differential responses to Phage-Antibiotic treatment

Since the phage-antibiotic treatment was only tested on a single bacterial isolate, we then aimed to test the efficacy of our phage-antibiotic cocktail on a panel of MDR strains of *S. marcescens*. The panel was composed of UT-383^30^, ten strains were obtained from clinical isolates from the CDC Antimicrobial Resistance Isolate Bank^31^, and three isolates were obtained from the University of Maryland from soil or plant rhizosphere samples from regional farms^32^. The MIC and susceptibility patterns for ampicillin, gentamicin, tetracycline, ciprofloxacin, and trimethoprim for each isolate is shown in **Table 2**. We observed that several of the CDC isolates (i.e. AR0517, AR0521 and AR0608) along with UT-383 had a broader resistance profile than all other tested isolates. When testing the differential susceptibility profiles of ampicillin, gentamicin, tetracycline, ciprofloxacin, and trimethoprim on all 14 non-MDR isolates via unsupervised hierarchical clustering, we observed that there were 2 different clusters defined by MIC (**Table 2**, **Fig. 5A**). The first cluster was defined as “MDR Plus” consists of CDC isolates AR0517, AR0521 and AR0608, along with UT-383 and the second cluster was defined as “MDR Base” showing high MIC for at least 3 antibiotics. To establish a baseline for the growth kinetics of all 14 isolates, we generated growth curves for each isolate for up to 40 hours (**Fig. 5B**). The data revealed that most of the isolates had a similar growth rate except for AR0027, which showed lower logarithmic growth in THB media (**Fig. 5B**). When testing the activity of our phage-antibiotic cocktail on the isolates, we challenged planktonic cultures after 12 hours in late Log phase at a concentration of 10 µg/mL of the antibiotic mixture combined with MOI 0.1 of the phage (**Fig. 5C-D**). The MDR Base isolates showed high susceptibility to the phage-antibiotic cocktail with a maintained reduction in absorbance 3 hours post treatment. However, the MDR Plus isolates showed tolerance to the phage-antibiotic cocktail. These observations underscore the effectiveness and limitations of phage-antibiotic mixtures in overcoming MDR.

**Figure 5:**
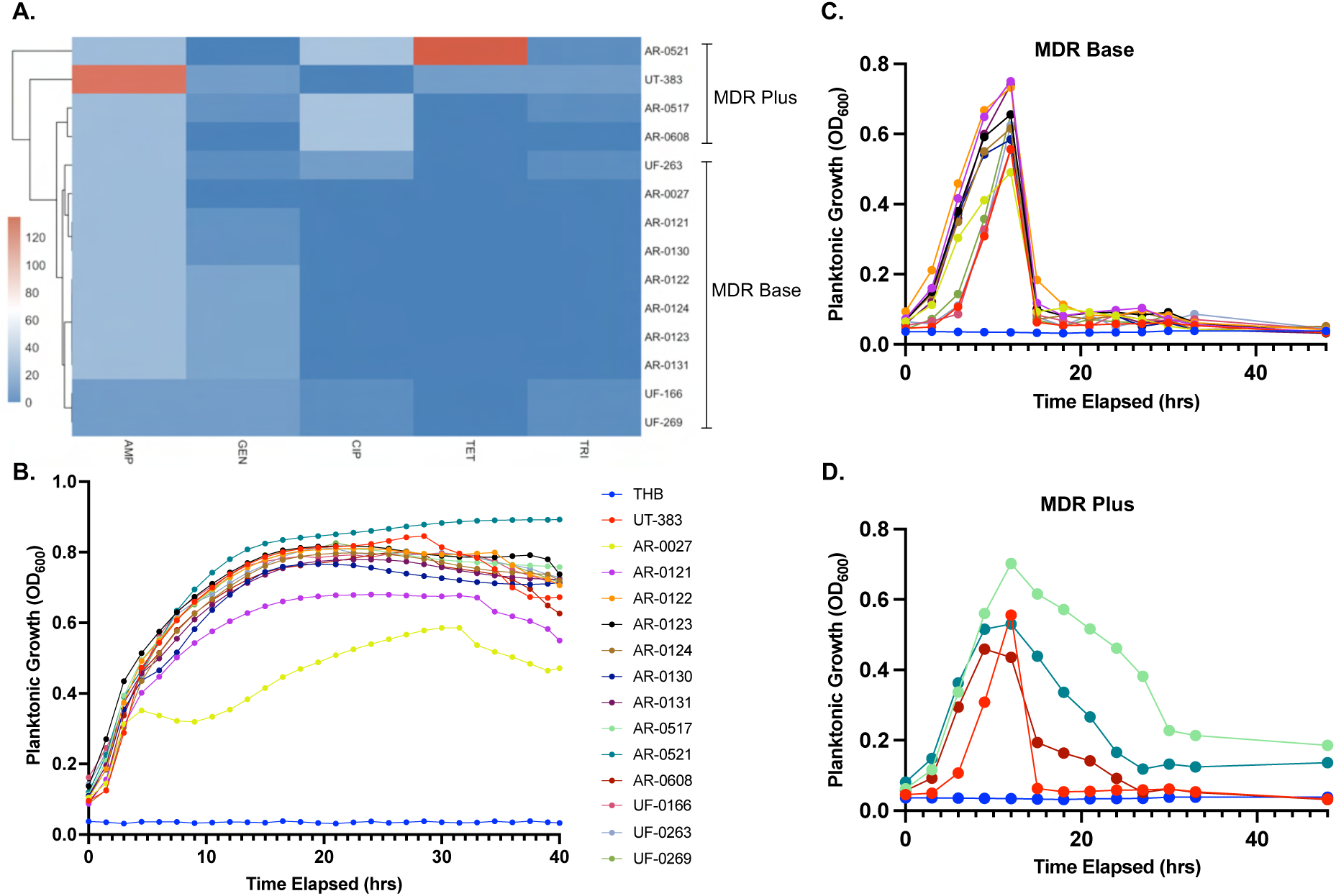
Phage-antibiotic combination therapy reduces growth of multi- and pan-drug resistant planktonic *S. marcescens*. **A**. Heat map comparing the minimum inhibitory concentrations (MICs) of various antibiotics across the 10 multidrug-resistant *Serratia marcescens* strains obtained from the CDC as well as 3 strains isolated from Maryland farms. **B.** OD_600_ planktonic growth curve data for the 14 tested strains measured over 40 hours without antibiotics. **C**. Growth curve conducted with the MDR Base from panel A challenged with the phage-antibiotics cocktail. **D**. Growth curve conducted with the MDR Plus strains from panel A. challenged with the phage-antibiotic cocktail.

### A triple combination of sub-MIC antibiotics, bacteriophages, and antimicrobial peptides eradicates biofilms and overcomes resistance in tolerant strains

Since the combination of phage and antibiotics was insufficient to eradicate all strains of *S. marcescens* tested, we questioned if integration of antimicrobial peptides (AMPs) active against a wide range of Enterobacterales, previously isolated by our group from *Lentilactobacillus hilgardii*^33^ into our phage-antibiotic cocktail could enhance treatment efficacy against mature *S. marcescens* biofilms. Mechanistically, this combined BAP approach leverages the combined actions of Penicillin-Streptomycin disrupting peptidoglycan synthesis and protein translation, Kanamycin halting protein production via 30S ribosomal binding, Ciprofloxacin impairing DNA gyrase and topoisomerase IV, AMPs presumably compromising membrane integrity through pore formation, and bacteriophages inducing bacterial replication hijacking and/or lysis (**Fig. 6A**). Disk diffusion assays demonstrated that in MDR Base strains, the BAP treatment produced the largest inhibition zones, marginally surpassing the phage-antibiotic combination and exceeding antibiotics alone (**Fig. 6B**). In MDR Plus isolates, inhibition zones were significantly reduced across single and double treatments. However, the BAP therapy achieved a mean zone of 4.43 mm, a 15.73-fold increase over the THB control (**Fig. 6C**), highlighting the potency of our combinatorial approach even against MDR Plus isolates.

**Figure 6:**
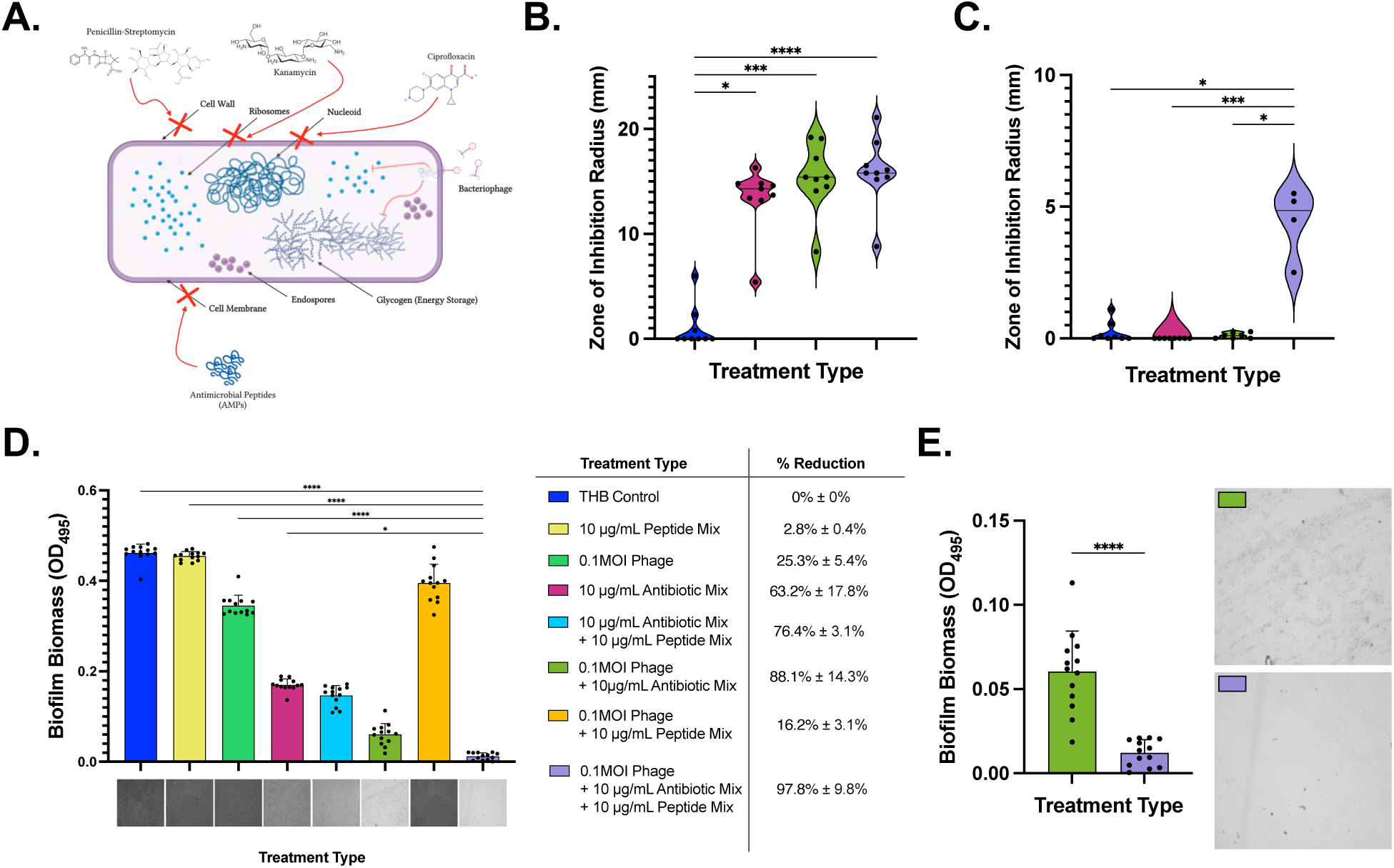
Synergistic combination of BAP achieves complete decolonization of *S. marcescens* biofilms. **A**. Schematic of antimicrobial targets of phages, antibiotics and peptides. Zone of inhibition analysis using disk diffusion assays of bacteriophages ϕJCVI_SM1 + ϕJCVI_SM2, antibiotics and peptides in all permutations against mature biofilms of **B**. *S. marcescens* MDR isolates, or **C.** *S. marcescens* PDR isolates. **D**. Biofilm biomass quantification after crystal violet staining (490 nm) of *S. marcescens* following treatment with antibiotics, bacteriophage, peptides, and permutations thereof. **E.** Biofilm biomass of the Phage + antibiotics mix compared to BAP cocktail. Statistical differences were determined by the Kruskal–Wallis test with Dunn’s multiple-comparison post-test. * p ≤ 0.05, ** p ≤ 0.01, *** p ≤ 0.001, **** p ≤ 0.0001. Error bars denote standard deviation.

Next, we tested BAP against mature biofilms of *S. marcescens* AR-0517, the CDC isolate with the highest observed antibiotic resistance profile. Absorbance quantification confirmed that the triple antimicrobial treatment reduced biomass by 97.8% relative to the untreated control (**Fig. 6D-E**). This surpassed the reductions observed with antibiotics alone (63.2%), phage + antibiotics (88.1%), antibiotics + peptides (76.4%), and phage + peptides (16.2%) permutations, while the peptides alone showed negligible anti-biofilm activity (**Fig. 6D**). Microscopic evaluation of mature *S. marcescens* biofilms corroborated these findings, showing complete biofilm eradication with the BAP treatment (**Fig. 6D-E**). Viability assays confirmed that biomass reductions corresponded to bacterial killing, with the BAP treatment achieving near-total clearance (>99.99% reduction) in Log CFU/mL in both supernatant and biofilm fractions compared to untreated controls (**Fig. 7A-C**). To support these results, we stained BAP-treated and untreated biofilms with a live-dead stain and observed that there was a stark reduction in green (live) fluorescence in the treated biofilms, with BAP treatment being the most effective. Mean fluorescence intensity (MFI) analyses of live-stained cells revealed a stepwise decline with treatment intensification (**Fig. 7D**). Together, this data confirms that biomass loss corresponds to reductions in bacterial viability, not just matrix disruption and well-clearance.

**Figure 7:**
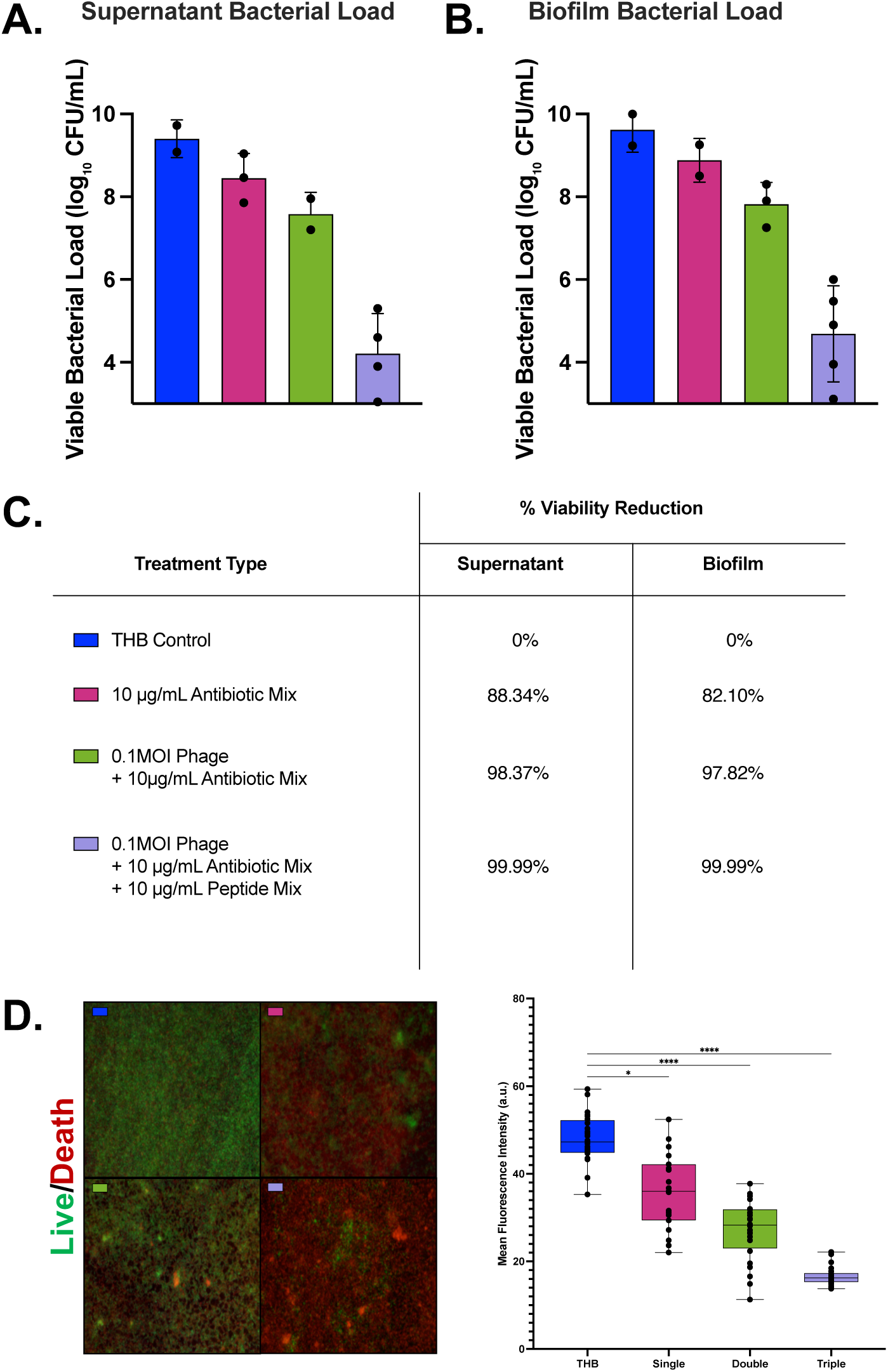
Synergistic combination of BAP achieves complete eradication of *S. marcescens* biofilms. CFU of *S. marcescens* biofilm **A**. supernatant or **B**. plate bound bacteria after treatment with the antibiotic mix, the phage + antibiotics mix and the BAP cocktail. **C**. Normalized data analysis of CFU reductions. **D**. Live/death fluorescent staining and mean fluorescent intensity measurement for the Live (green) and Death (red) channels. Statistical differences were determined by the Kruskal–Wallis test with Dunn’s multiple-comparison post-test. * p ≤ 0.05, ** p ≤ 0.01, *** p ≤ 0.001, **** p ≤ 0.0001. Error bars denote standard deviation.

Finally, *S. marcescens* is known to cause severe infections associated with the respiratory tract^30,34,35^. We therefore evaluated the toxicity of BAP in human airway epithelial cells by means of a Lactate Dehydrogenase (LDH) assay. No observable toxicity was detected after 12- or 24-hours post-treatment (**Supplemental Fig. 2**). Collectively, these results demonstrate that the triple antimicrobial BAP combination strategy effectively breaks through *S. marcescens* biofilm defense and resistance mechanisms, achieving superior biofilm disruption and bacterial clearance in MDR strains without harming mammalian cells.

## Discussion

The management of *S. marcescens* infections continues to be hindered by its intrinsic and biofilm-associated resistance to a wide range of antibiotics. Here, we demonstrate that multidrug-resistant *S. marcescens* sourced from various environments exhibits variable susceptibility to distinct antibiotic classes depending on its physical state, as planktonic cells or in biofilm. Initial screening using antibiotics that target major bacterial processes, including cell wall synthesis (β-lactams), protein synthesis (aminoglycosides, tetracyclines, chloramphenicol), DNA replication (fluoroquinolones), and folate metabolism (sulfonamides), revealed significant differences in their efficacy which may be due to *S. marcescens* broad genomic diversity^36^. While several agents, such as Penicillin-Streptomycin, Ciprofloxacin, and Kanamycin, substantially inhibited planktonic bacteria at high concentrations, their effects as single therapeutics were markedly diminished on mature biofilms. Among the tested agents, Penicillin-Streptomycin, Kanamycin and Ciprofloxacin retained some biofilm-disruptive capacity, while others like Gentamicin and Trimethoprim showed only partial effects. These results highlight the limitations of monotherapy and underscore the therapeutic challenge posed by biofilm-associated infections^17^. Importantly, we then showed that combining antibiotics targeting distinct cellular pathways leads to enhanced clearance of both planktonic and biofilm-associated *S. marcescens*, and that this effect is further amplified when bacteriophages and antimicrobial peptides are added. This multi-tier antimicrobial combination was observed to eradicate 99.99% of mature *S. marcescens* biofilms. Together, these findings emphasize the importance of multidimensional antimicrobial strategies for overcoming the resistance of *S. marcescens* biofilms in community and nosocomial environments.

For *S. marcescens* and other nosocomial isolates, susceptibility is generally highest for aminoglycosides (e.g. gentamicin, amikacin), third-generation cephalosporins (e.g. cefotaxime), fluoroquinolones (e.g. ciprofloxacin, levofloxacin), and trimethoprim-sulfamethoxazole (cotrimoxazole), although resistance trends are escalating^37,38^. Intrinsic resistance stems from chromosomally encoded AmpC β-lactamases and efflux pumps like SdeAB, which confer resistance to penicillins, first- and second-generation cephalosporins, chloramphenicol, fluoroquinolones, and tetracyclines^39^. Acquired resistance occurs most commonly through plasmid horizontal gene transfer or through genetic mutations^7^. In the context of biofilms, high-dose antibiotic lock therapy with chloramphenicol has demonstrated significant biofilm biomass reduction and bacterial viability suppression on catheter-like surfaces *in vitro*^8^. However, chloramphenicol efficacy can be limited by efflux pump-mediated resistance pathways^40^. Similarly, combination strategies using antibiotic-loaded mesoporous silica nanoparticles (MSNs) have shown near-complete eradication of biofilms when antibiotics such as levofloxacin or gentamicin are stably released over prolonged periods, although direct application to *S. marcescens* remains to be tested^41^. Thus, to date, there is limited knowledge of how new antibiotic mono- or combinatorial strategies may be effective against *S. marcescens* biofilms.

Bacteriophages have emerged as promising therapeutic alternatives for combating AMR pathogens, including *S. marcescens*. Phage therapy offers pathogen-specific lytic activity, the potential to penetrate biofilms, and minimal impact on commensal microbiota, making it a particularly attractive option for multidrug-resistant *S. marcescens* infections. Studies have identified and characterized lytic phages active against clinical *S. marcescens* isolates, demonstrating differential efficacy in both planktonic and biofilm-associated states^42,43^. Moreover, phage-antibiotic synergy has been observed for other antimicrobial-resistant pathogens such as *Acinetobacter baumannii* and *Pseudomonas aeruginosa*, suggesting that combination therapy could enhance bacterial clearance and prevent resistance development^44^. However, no such study has been conducted for *S. marcescens* biofilms.

As resistance to conventional antibiotics continues to rise, especially among nosocomial pathogens, like *S. marcescens*, multi-tiered antimicrobial interventions represent a viable adjunct or alternative strategy warranting further investigation in both preclinical and clinical settings. We also show that in addition to antibiotics and bacteriophages, the integration of antimicrobial peptides provided a synergistic approach to promote complete biofilm decolonization. Antimicrobial peptides are short, positively charged, amphipathic molecules that can have bactericidal activity mainly through membrane permeabilization^45^. Other mechanisms associated with AMPs are the targeting of intracellular bacterial processes and host immune modulation^45^. When combined with bacteriophages and/or conventional antibiotics, AMPs can promote the reduction of antibiotic resistance and lead to synergistic killing effects^46,47^. Thus, our results provide a framework to use multi-tier antimicrobial strategies for the full decolonization of *S. marcescens* biofilms and its translation to other nosocomial pathogens.

In conclusion, the multifactorial aspect of bacterial antibiotic resistance mechanisms requires innovative experimental approaches that factor these lines of defense. Our proposed approach provides key experimental data with translational potential. Future studies should target the therapeutic potential of multi-tier antimicrobial therapies in infection models *in vitro* and *in vivo* as well as a detailed understanding of the synergistic mechanism of action.

## Methods

### Planktonic Growth and Quantification

*Serratia marcescens* cultures were struck onto Blood Agar plates from prepared stock and incubated overnight. A single colony was grown in 4 mL Todd Hewitt Broth (THB) (Sigma-Aldrich, USA) at 37°C for 24 hours until the culture reached an optical density of 1.24 (OD_600_) with an equivalence to ∼10^8^ CFU/mL. After 24 hours of incubation, the culture was diluted with 4 mL of fresh THB media to a total of 8 mL with an approximate OD_600_ of 0.6 to start the growth curve culture. Growth measurements were recorded spectrophotometrically at OD_600_ every 30 minutes for 48 hours. Growth curves for the 14 strains of *S. marcescens* were generated in a 96 well format using the Multiskan FC spectrophotometer (ThermoFisher, Waltham, MA) and measuring the absorbance (OD_490_) or optical density (OD_600_) every 30-90 minutes for 24-48 hours.

### Biofilm Growth, Staining and Quantification

Biofilms were established by streaking *S. marcescens* liquid cultures onto blood agar plates and incubating them at 37°C overnight to obtain isolated single colonies. A single colony was inoculated into 4 mL of THB medium (Sigma-Aldrich, USA) and incubated at 37°C for 4-8 hours. After reaching an OD_600_ of 1.20, cultures were diluted by adding 12 mL of fresh THB media and transferred to well plates in the following volumes: 1 mL total (200 µL culture + 800 µL THB) for 24-well plates, 500 µL total (100 µL culture + 400 µL THB) for 48-well plates, and 250 µL total (50 µL culture + 200 µL THB) for 96-well plates. Plates were incubated statically at 37°C for 24 hours to allow biofilm formation. At 24 hours, the media was gently aspirated and replaced with fresh THB media before incubating the plates for an additional 24 hours. To quantify the biofilms, media was removed, washed 3 times with sterile phosphate buffered saline (PBS) (Gibco, pH 7.4) and air-dried, then fixed with 4% formaldehyde (Sigma-Aldrich, USA) for 10 minutes. After fixation, formaldehyde was removed, and biofilms were stained with 1% crystal violet for 10-15 minutes. Excess stain was removed, and wells were washed three times with PBS. Biofilms were then air-dried for 5-10 minutes before imaging on a Leica DMi1 microscope. To quantify biofilm biomass, 200 µL of 70% ethanol was added to each well to extract the crystal violet stain, and absorbance was measured at OD_490_ using a Varioskan Lux microplate reader (Thermo Fisher Scientific, Waltham, MA).

### Phage Wastewater Processing

Bacteriophages were isolated from untreated inflow wastewater obtained from a local water treatment facility in Poolesville, Maryland, USA. Samples were first centrifuged in 100 mL aliquots for 30 minutes at 7,500×g, at 4°C to pellet solids and sediment. Supernatants were then vacuum filtered through a 0.2 µm filter to remove bacteria and stored for future experiments.

### Phage Enrichment

*S. marcescens* bacterial isolates UT-383^30^ and HER1170^48^ were inoculated in Tryptic Soy broth (TSB) (Sigma-Aldrich, USA) and grown overnight at 37°C shaking. The next day, 20 mL of fresh TSB, 5 mL of processed wastewater, and 50 µL of overnight host bacteria culture were mixed in a 150mL Erlenmeyer flask and again incubated overnight at 37°C shaking. The next day “enriched” cultures were transferred to 50 mL conical centrifuge tubes and spun at 8,000 × g for 15 minutes to pellet bacteria cells. Supernatants were then vacuum filtered through a 0.2 µm filter to remove bacteria.

### Isolation of Bacteriophages

Bacteriophages were identified using a soft agar overlay assay. Briefly, 150 mm bottom agar petri plates were prepared containing TSB with 1.5% agar, 1 mM CaCl_2_ and 1 mM MgCl_2_. *S. marcescens* bacterial isolates MB383 and HER1170 were inoculated into TSB and grown overnight at 37°C. 300 µL of stationary phase culture and 300 µL of “enriched” wastewater were added to a tube and incubated at room temp for 10 minutes. Then, 10 mL of top agar (TSB with 0.7% agarose, 1 mM CaCl_2_, and 1 mM MgCl_2_) cooled to 55°C was added and the mixture was plated onto a bottom agar plate and solidified. Plates containing bacteria culture and top agar only were used as a negative control. Plates were incubated overnight at 37°C and were examined the following day to identify lytic phage plaques. Phages were passaged by sequential picking and plating of individual plaques grown on the same bacterial isolate. Phages were picked from individual plaques using a pipette tip and were placed into 100 μL of SM buffer (50 mM Tris-HCl pH 7.5, 100 mM NaCl, 8 mM MgSO_4_) and incubated overnight at 37°C. The following day, 10-fold serial dilutions were made in SM buffer, and 10 μL of each dilution was spotted onto a plate containing 5 mL of top agarose mixed with 100 μL of stationary phase bacterial culture and layered on top of bottom agar. After overnight incubation at 37°C, an individual plaque was picked and passaged again. Each phage was passaged three times before the generation of high-titer stocks.

### Phage Stock Generation

To generate high-titer stocks, a single plaque was picked into 100 μL of SM buffer and incubated overnight. Then, 100 μL of overnight bacterial culture (OD_600_ > 1.0) was added and the mixture was incubated for 5-10 min at room temperature, followed by addition of 10 mL of top agarose and plating onto large bottom agar plates. Plates were incubated overnight at 37°C and then plates with high plaque density were flooded with 10 mL of SM buffer and incubated at 37°C for at least 1 h to elute phages from the top agar. The SM buffer was removed from each plate, pooled, spun down at 4,000 rpm for 20 min, and filtered through a 0.2 μm filter, titered, and stored at 4°C.

### Peptide Synthesis

The four bioinformatically-predicted peptide sequences from *Lentilactobacillus hilgardii* previously published by our group were chemically synthesized by LifeTein LLC to a minimum of 85% purity^33^. The working concentrations spanned from 5 to 15 μg/mL.

### Bacteriophage, Antibiotics and Peptides (BAP) Treatment

Both planktonic and biofilm cultures were treated with BAP. Antibiotic solutions were diluted in THB media from their starting concentrations to the intended doses (0.8 mg/mL and 0.08 mg/mL)^8^. The combination antibiotic treatment consisted of equal concentrations of penicillin-streptomycin, gentamicin, and chloramphenicol at a dose of 33.3 µg/mL each, totaling 100 µg/mL for our starting treatment. Bacteriophages ϕJCVI_Sm1 and ϕJCVI_Sm2 were diluted in SM buffer to multiplicity of infection (MOI) of 0.01, 0.1, 1, 10, and 100. Peptides were diluted from stock concentrations (40 μg/mL) in molecular biology grade water (Corning, Manassas, VA) to a final concentration of 10 μg/mL.

Planktonic cultures were grown as indicated above and 250 µL of culture media was added into each well of a 24-well plate. An additional 250 µL of 0.8 mg/mL and 0.08 mg/mL of antibiotics, 10 μg/mL concentration of peptides and a respective MOI (selected from a range of 0.01 to 100 in a separate experiment) of bacteriophage solution were added to each well for a total of 500 µL of media per well. The plates were then incubated at 37°C for 24 hours with shaking at 120 rpm and optical density readings were taken in increments of 30 minutes for 48 hours at 37°C with shaking at 120 rpm to assess the effects of the treatment on bacterial growth dynamics.

For biofilms, the cultures were grown as indicated above until right before the fixation and quantification. Instead of fixation, the media in the wells was aspirated and the biofilms received 250mL of THB media along with 250 µL of 0.8 mg/mL and 0.08 mg/mL of antibiotics, 10 μg/mL concentration of peptides and a respective MOI of bacteriophage solution per well. Then the biofilms were incubated for another 24 hours at 37°C without shaking to allow for proper and thorough treatment. After this 24-hour incubation, biofilms were aspirated again and washed with sterile PBS before the staining, imaging and quantification were done as described above.

### Disk Diffusion Assay

To evaluate the zone of inhibition induced by different treatment combinations, a disk diffusion assay was performed using all 14 *S. marcescens* strains described in this study. Each strain was cultured in 4 mL Todd Hewitt Broth (THB) at 37°C for 24 hours until the culture reached an optical density around 1.3 (OD_600_) with an equivalence to ∼10^8^ CFU/mL. Then 100 μL of the solution was uniformly spread using a sterile cotton swab to ensure even bacterial lawn formation. For each strain, the plate was divided into 4 quadrants, and four sterile paper disks (Microxpress, Goa, India) were placed equidistantly, one on each quadrant on the agar surface. The first disk served as a negative control, containing only Todd Hewitt Broth (THB) without any antimicrobial agents. The second disk contained the cocktail antibiotic solution at 100 µg/mL, composed of penicillin-streptomycin, gentamicin, and chloramphenicol. The third disk consisted of the same antibiotic combination supplemented with bacteriophage at an MOI of 0.1. The fourth disk contained the full treatment, which consisted of the antibiotic cocktail, bacteriophage (MOI 0.1), and 10 µg/mL of AMPs from *L. hilgardii*. Plates were incubated at 37°C for 24 hours, after which the diameter of the inhibition zones surrounding each disk was measured using a ruler. These measurements allowed for a comparative assessment of antimicrobial efficacy across treatment types and strains, with larger zones indicating higher levels of bacterial growth inhibition.

### Minimum Inhibitory Concentration Assay

An overnight culture of *S. marcescens* was started and was diluted 1:10,000 the next day in fresh THB media. All antibiotics used were sterile filtered for this assay and used at the indicated concentrations. First, serial two-fold dilutions of antibiotics were prepared starting at the highest concentration. Then, 100µl of antibiotic at the indicated concentration was mixed with 100µl of diluted *S. marcescens* culture in each well of a 96 well plate, in triplicate. The plate was then incubated at 37°C overnight and was observed the next day for the lowest concentration of antibiotic that inhibited bacterial growth. This concentration was determined to be the minimum inhibitory concentration.

### Colony Forming Unit Viability Assay

We performed Colony Forming Unit (CFU) viability assays following treatment with our BAP mixture to assess its bactericidal effect on *S. marcescens*. After treatment, cultures were separated into the supernatant and the scraped biofilm fraction. Each was serially diluted in sterile PBS, and 10 µL of each dilution was spot-plated onto blood agar plates.

### Mean Fluorescence Intensity Imaging

*S. marcescens* biofilms were grown overnight in 200 µL of THB at 37°C in an 8 Well Chamber, removable slide (Ibidi, cat. 80841), then exposed for 24 hours to THB alone (control) or to our Single, Double, or Triple treatment. After aspiration of the supernatant, biofilms were stained using the LIVE/DEAD BacLight Bacterial Viability kit (Invitrogen, USA) following manufacturers protocol. Fluorescence images were captured immediately on a Leica SP8 confocal microscope (4x objective). For each well, five non-overlapping Z-stacks (20 µm, 1 µm steps) were acquired and rendered as maximum-intensity projections. Mean fluorescence intensity for green (live) and red (dead) channels was quantified in Fiji after background subtraction and averaged across fields. Images included in the manuscript were processed in ImageJ (version 1.54r 25) to separate the fluorescent channels (red and green).

### Lactate Dehydrogenase release assay

The Lactate Dehydrogenase (LDH) assay was used to test the toxicity of the combination of sub-MIC antibiotics, peptides and phages in alveolar type II epithelial cells (A549, ATCC CCL-185). Briefly, A549 cells were maintained in sterile-filtered DMEM supplemented with 10% FBS and 5% CO₂. For LDH assay, cells were seeded in 96-well plates at 5 × 10⁴ cells/well.

Media was replaced with RPMI 1640 containing 2% FBS, and the cocktail of sub-MIC antibiotics, peptides and phages. Plates were incubated for 12 or 24 hours at 37 °C, 5% CO₂. LDH release was quantified using the Invitrogen™ CyQUANT LDH Cytotoxicity Assay Kit per the manufacturer’s instructions. Absorbance was measured at OD_490_ and OD_680_ using the Varioskan LUX (Thermo Fisher, Waltham, MA). Data analysis was conducted as mentioned below. Experiments were performed in triplicate.

### Statistical Analysis

All experiments were conducted with 8-24 biological replicates per strain and corresponding controls to ensure statistical significance. Kruskal-Wallis tests along with Dunn’s multiple comparisons testing were performed using GraphPad Prism version 10.4.1 to assess the significance of the differences between planktonic and biofilm growth in response to antibiotic, bacteriophage, and peptide treatments. For select pairwise comparisons where we wanted to emphasize the effect of specific treatments, Mann-Whitney U tests were also conducted to evaluate significance more directly. For the heat map representing treatment responses across strains, we performed a hierarchical cluster analysis to group strains based on their resistance profiles and phenotypic similarities. Lastly, to determine statistical significance for the MFI quantification, the biological replicates per treatment were analyzed using one-way ANOVA with Tukey’s post-test (α = 0.05). Significance was determined at p < 0.05 (*), p < 0.01 (**), p < 0.001 (***), and p < 0.0001 (****).

## Acknowledgements

We thank the CDC Antimicrobial Resistance Isolate Bank for providing *Serratia marcescens* clinical isolates. The opinions, assertions and contents are solely the responsibility of the authors and do not necessarily represent the official views of CDC.

## Funding Information

This research was funded by cooperative agreement U54CK000603 (DEF) from the United States Centers for Disease Control and Prevention (CDC), by R01 AI168313 (NG-J) from the National Institutes of Health National Institute of Allergy and Infectious Diseases and by the University of Maryland College Park Start-up Funds (NG-J).

## Author Contributions

Conceptualization: NG-J, AD. Data curation: NG-J, AD, AA, IV. Formal analysis: NG-J, AD, IV. Funding acquisition: NG-J, DF. Investigation: NG-J, AD, AA, IV, HN, EH, NF, KT. Methodology: NG-J, AD, HN, EH. Project administration: NG-J. Resources: NG-J, DF, and RB. Software: N/A. Supervision: NG-J. Validation: NG-J, AD, AA, IV, HN, EH, NF, KT. Visualization: NG-J, AD, AA, HN. Writing – original draft: NG-J, AD. Writing – review & editing: NG-J, AD, AA, DF, RB.

## Conflict of Interest Statement

The authors declare no conflicts of interest.

## Notes

### Competing Interest Statement

The authors have declared no competing interest.

## References

1 Hejazi, A. & Falkiner, F. R. Serratia marcescens. J Med Microbiol 46, 903–912 (1997). 10.1099/00222615-46-11-903

2 Redondo-Bravo, L. et al. Serratia marcescens outbreak in a neonatology unit of a Spanish tertiary hospital: Risk factors and control measures. Am J Infect Control 47, 271–279 (2019). 10.1016/j.ajic.2018.08.026

3 Mendes, J. C. & Casado, A. Serratia marcescens outbreak in a COVID-19 intensive care unit - Are there any factors specific to COVID-19 units that facilitate bacterial cross-contamination between COVID-19 patients? Am J Infect Control 50, 223–225 (2022). 10.1016/j.ajic.2021.10.005

4 Kim, E. J. et al. Outbreak investigation of Serratia marcescens neurosurgical site infections associated with a contaminated shaving razors. Antimicrob Resist Infect Control 9, 64 (2020). 10.1186/s13756-020-00725-6

5 Suetens, C. et al. Prevalence of healthcare-associated infections, estimated incidence and composite antimicrobial resistance index in acute care hospitals and long-term care facilities: results from two European point prevalence surveys, 2016 to 2017. Euro Surveill 23 (2018). 10.2807/1560-7917.ES.2018.23.46.1800516

6 Ramasethu, J. Prevention and treatment of neonatal nosocomial infections. Matern Health Neonatol Perinatol 3, 5 (2017). 10.1186/s40748-017-0043-3

7 Tavares-Carreon, F., De Anda-Mora, K., Rojas-Barrera, I. C. & Andrade, A. Serratia marcescens antibiotic resistance mechanisms of an opportunistic pathogen: a literature review. PeerJ 11, e14399 (2023). 10.7717/peerj.14399

8 Ray, C., Shenoy, A. T., Orihuela, C. J. & Gonzalez-Juarbe, N. Killing of Serratia marcescens biofilms with chloramphenicol. Ann Clin Microbiol Antimicrob 16, 19 (2017). 10.1186/s12941-017-0192-2

9 Yang, H. et al. Discovery of a fluoroquinolone-resistant Serratia marcescens clinical isolate without quinolone resistance-determining region mutations. Ann Lab Med 34, 487–488 (2014). 10.3343/alm.2014.34.6.487

10 Cosimato, I. et al. Current Epidemiological Status and Antibiotic Resistance Profile of Serratia marcescens. Antibiotics (Basel) 13 (2024). 10.3390/antibiotics13040323

11 Bryant, K. A. et al. KPC-4 Is encoded within a truncated Tn4401 in an IncL/M plasmid, pNE1280, isolated from Enterobacter cloacae and Serratia marcescens. Antimicrob Agents Chemother 57, 37–41 (2013). 10.1128/AAC.01062-12

12 Farrar, W. E., Jr. & O’Dell N, M. beta-Lactamases and resistance to penicillins and cephalosporins in Serratia marcescens. J Infect Dis 134, 245–251 (1976). 10.1093/infdis/134.3.245

13 Flemming, H. C. et al. Biofilms: an emergent form of bacterial life. Nat Rev Microbiol 14, 563–575 (2016). 10.1038/nrmicro.2016.94

14 Donlan, R. M. Biofilms and device-associated infections. Emerg Infect Dis 7, 277–281 (2001). 10.3201/eid0702.010226

15 Donlan, R. M. Biofilm formation: a clinically relevant microbiological process. Clin Infect Dis 33, 1387–1392 (2001). 10.1086/322972

16 Buzzo, J. R. et al. Z-form extracellular DNA is a structural component of the bacterial biofilm matrix. Cell 184, 5740–5758 e5717 (2021). 10.1016/j.cell.2021.10.010

17 Hall, C. W. & Mah, T. F. Molecular mechanisms of biofilm-based antibiotic resistance and tolerance in pathogenic bacteria. FEMS Microbiol Rev 41, 276–301 (2017). 10.1093/femsre/fux010

18 Zivkovic Zaric, R., et al. Antimicrobial Treatment of Serratia marcescens Invasive Infections: Systematic Review. Antibiotics (Basel) 12 (2023). 10.3390/antibiotics12020367

19 Bourigault, C. et al. Duodenoscopy: an amplifier of cross-transmission during a carbapenemase-producing Enterobacteriaceae outbreak in a gastroenterology pathway. J Hosp Infect 99, 422–426 (2018). 10.1016/j.jhin.2018.04.015

20 Oie, S., Yoshida, H. & Kamiya, A. Microbial contamination of water-soaked cotton gauze and its cause. Microbios 104, 159–166 (2001).

21 Hanczvikkel, A. et al. Nosocomial outbreak caused by disinfectant-resistant Serratia marcescens in an adult intensive care unit, Hungary, February to March 2022. Euro Surveill 29 (2024). 10.2807/1560-7917.ES.2024.29.26.2300492

22 Wang, X., Fan, F., Dong, S. & Zhang, Y. Emergence of carbapenem-resistant Serratia marcescens co-harboring bla(NDM-1), bla(KPC-2), and bla(SRT-2) in bloodstream infection. Microbiol Spectr 13, e0054525 (2025). 10.1128/spectrum.00545-25

23 Stewart, P. S., Rayner, J., Roe, F. & Rees, W. M. Biofilm penetration and disinfection efficacy of alkaline hypochlorite and chlorosulfamates. J Appl Microbiol 91, 525–532 (2001). 10.1046/j.1365-2672.2001.01413.x

24 McCarlie, S. & Bragg, R. R. Impact of the Stress Response on Quaternary Ammonium Compound Disinfectant Susceptibility in Serratia Species. Microorganisms 12 (2024). 10.3390/microorganisms12112240

25 Pal, C., Bengtsson-Palme, J., Kristiansson, E. & Larsson, D. G. Co-occurrence of resistance genes to antibiotics, biocides and metals reveals novel insights into their co-selection potential. BMC Genomics 16, 964 (2015). 10.1186/s12864-015-2153-5

26 Ayub, A., Cheong, Y. K., Castro, J. C., Cumberlege, O. & Chrysanthou, A. Use of Hydrogen Peroxide Vapour for Microbiological Disinfection in Hospital Environments: A Review. Bioengineering (Basel) 11 (2024). 10.3390/bioengineering11030205

27 Bourdin, T. et al. Disinfection of sink drains to reduce a source of three opportunistic pathogens, during Serratia marcescens clusters in a neonatal intensive care unit. PLoS One 19, e0304378 (2024). 10.1371/journal.pone.0304378

28 Fernandez-Hidalgo, N. & Almirante, B. Antibiotic-lock therapy: a clinical viewpoint. Expert Rev Anti Infect Ther 12, 117–129 (2014). 10.1586/14787210.2014.863148

29 Kapoor, G., Saigal, S. & Elongavan, A. Action and resistance mechanisms of antibiotics: A guide for clinicians. J Anaesthesiol Clin Pharmacol 33, 300–305 (2017). 10.4103/joacp.JOACP_349_15

30 Gonzalez-Juarbe, N. et al. Requirement for Serratia marcescens cytolysin in a murine model of hemorrhagic pneumonia. Infect Immun 83, 614–624 (2015). 10.1128/IAI.01822-14

31 Lutgring, J. D. et al. FDA-CDC Antimicrobial Resistance Isolate Bank: a Publicly Available Resource To Support Research, Development, and Regulatory Requirements. J Clin Microbiol 56 (2018). 10.1128/JCM.01415-17

32 Harrelson, E., Zeng, Q., Gao, M., Toro, M. & Blaustein, R. A. Multidrug resistance in bacteria associated with leafy greens and soil in urban agriculture systems. Front Plant Sci 16, 1664284 (2025). 10.3389/fpls.2025.1664284

33 Appel, A. et al. In Silico Identification and Molecular Characterization of Lentilactobacillus hilgardii Antimicrobial Peptides with Activity Against Carbapenem-Resistant Acinetobacter baumannii. Antibiotics (Basel) 14 (2025). 10.3390/antibiotics14101004

34 Gonzalez-Juarbe, N. et al. Pore-Forming Toxins Induce Macrophage Necroptosis during Acute Bacterial Pneumonia. PLoS Pathog 11, e1005337 (2015). 10.1371/journal.ppat.1005337

35 Gonzalez-Juarbe, N. et al. Pore-forming toxin-mediated ion dysregulation leads to death receptor-independent necroptosis of lung epithelial cells during bacterial pneumonia. Cell Death Differ 24, 917–928 (2017). 10.1038/cdd.2017.49

36 Abreo, E. & Altier, N. Pangenome of Serratia marcescens strains from nosocomial and environmental origins reveals different populations and the links between them. Sci Rep 9, 46 (2019). 10.1038/s41598-018-37118-0

37 Weber, L., Jansen, M., Kruttgen, A., Buhl, E. M. & Horz, H. P. Tackling Intrinsic Antibiotic Resistance in Serratia Marcescens with A Combination of Ampicillin/Sulbactam and Phage SALSA. Antibiotics (Basel) 9 (2020). 10.3390/antibiotics9070371

38 Khan, s. et al. in Exploring Bacterial Biofilms (eds Sadık Dincer, Melis Sumengen Ozdenefe, & Hatice Aysun Mercimek Takci) (IntechOpen, 2025).

39 Begic, S. & Worobec, E. A. The role of the Serratia marcescens SdeAB multidrug efflux pump and TolC homologue in fluoroquinolone resistance studied via gene-knockout mutagenesis. Microbiology 154, 454–461 (2008). 10.1099/mic.0.2007/012427-0

40 Vivekanandan, K. E., Kumar, P. V., Jaysree, R. C. & Rajeshwari, T. Exploring molecular mechanisms of drug resistance in bacteria and progressions in CRISPR/Cas9-based genome expurgation solutions. Glob Med Genet 12, 100042 (2025). 10.1016/j.gmg.2025.100042

41 Aguilar-Colomer, A. et al. Impact of the antibiotic-cargo from MSNs on Gram-positive and Gram-negative bacterial biofilms. Microporous Mesoporous Mater 311, 110681 (2021). 10.1016/j.micromeso.2020.110681

42 Carr, E. L. et al. Genome Sequences of 14 Siphophages That Infect Serratia marcescens. Microbiol Resour Announc 11, e0121221 (2022). 10.1128/mra.01212-21

43 Li, L. et al. Isolation, characterization, and genomic analysis of a novel phage WSPA with lytic activity against Serratia marcescens. Microbiol Spectr 13, e0271624 (2025). 10.1128/spectrum.02716-24

44 Morris, T. C., Reyneke, B., Khan, S. & Khan, W. Phage-antibiotic synergy to combat multidrug resistant strains of Gram-negative ESKAPE pathogens. Sci Rep 15, 17235 (2025). 10.1038/s41598-025-01489-y

45 Oliveira Junior, N. G., Souza, C. M., Buccini, D. F., Cardoso, M. H. & Franco, O. L. Antimicrobial peptides: structure, functions and translational applications. Nat Rev Microbiol 23, 687–700 (2025). 10.1038/s41579-025-01200-y

46 Taheri-Araghi, S. Synergistic action of antimicrobial peptides and antibiotics: current understanding and future directions. Front Microbiol 15, 1390765 (2024). 10.3389/fmicb.2024.1390765

47 Nayab, S., Idrees, K. & Aslam, M. A. Synergism of phages and antimicrobial peptides for treating multidrug resistant bacterial pathogens. Exploration of Drug Science 3, 1008133 (2025). 10.37349/eds.2025.1008133

48 Grimont, F. Les bactériophages des Serratia et bactéries voisines, Université de Bordeaux II, France., (1977).

