## Supplemental Tables and Figures for "A Triple-Modality Peptide-Antibiotic-Phage Therapy Eradicates Multidrug-Resistant *Serratia marcescens* Biofilms"

**Supplemental Figures and Tables:**

| Antibiotic | Mechanism of Action | Bacterial Component Targeted | Mode of Action |
|---|---|---|---|
| Ampicillin (Am) | Inhibits cell wall synthesis | Transpeptidase | Bactericidal |
| Chloramphenicol (Cm) | Inhibits protein synthesis | 50S ribosomal subunit | Bacteriostatic |
| Penicillin-Streptomycin (Pen-Strep) | Inhibits cell wall synthesis<br>Inhibits protein synthesis | Peptidoglycan synthesis<br>30S ribosomal subunit | Bactericidal |
| Tetracycline (Tc) | Inhibits protein synthesis | 30S ribosomal subunit | Bacteriostatic |
| Gentamicin (Gm) | Inhibits protein synthesis | 30S ribosomal subunit | Bactericidal |
| Kanamycin (Km) | Inhibits protein synthesis | 30S ribosomal subunit | Bactericidal |
| Ciprofloxacin (Cip) | Inhibits DNA replication | DNA gyrase<br>Topoisomerase II & IV | Bactericidal |
| Co-trimoxazole (Tri) | Inhibits folic acid synthesis | Dihydrofolate reductase<br>Dihydropteroate synthase | Bactericidal |

**Table 1: Bacterial targets of common antibiotics.**

| Isolate_ID | Organism Name | Ampicillin MIC | Gentamicin MIC | Tetracycline MIC | Ciprofloxacin MIC | Trimethoprim MIC |
|---|---|---|---|---|---|---|
| UT-383 | Serratia marcescens | >32 | 16 | 16 | 0.5 | 16 |
| AR-0027 | Serratia marcescens | >32 | ≤0.25 | 16 | 1 | 2 |
| AR-0091 | Serratia marcescens | >32 | 1 | 4 | ≤0.25 | ≤0.5 |
| AR-0099 | Serratia marcescens | >32 | 0.5 | 8 | 0.5 | ≤0.5 |
| AR-0121 | Serratia marcescens | >32 | 0.5 | 8 | ≤0.25 | ≤0.5 |
| AR-0122 | Serratia marcescens | >32 | 0.5 | 8 | ≤0.12 | ≤0.5 |
| AR-0123 | Serratia marcescens | >32 | 0.5 | 8 | ≤0.12 | ≤0.25 |
| AR-0124 | Serratia marcescens | >32 | 0.5 | 8 | ≤0.12 | 0.5 |
| AR-0130 | Serratia marcescens | >32 | 1 | >32 | 0.25 | ≤0.25 |
| AR-0131 | Serratia marcescens | >32 | 0.5 | 16 | 0.25 | ≤0.5 |
| AR-0517 | Serratia marcescens | >32 | >16 | >32 | 4 | >8 |
| AR-0520 | Serratia marcescens | >32 | 8 | 8 | 2 | >8 |
| AR-0521 | Serratia marcescens | >32 | >16 | >32 | >8 | >8 |
| AR-0608 | Serratia marcescens | >32 | 4 | 16 | 2 | ≤0.5 |
| UF-166 | Serratia marcescens | 16 | 16 | 8 | 0.25 | 8 |
| UF-263 | Serratia marcescens | 32 | 8 | 16 | 0.5 | 8 |
| UF-269 | Serratia marcescens | 16 | 16 | 8 | 0.5 | 8 |

| Isolate_ID | Organism Name | Ampicillin INT | Gentamicin INT | Tetracycline INT | Ciprofloxacin INT | Trimethoprim |
|---|---|---|---|---|---|---|
| UT-383 | Serratia marcescens | R | R | R | S | R |
| AR-0027 | Serratia marcescens | R | S | R | R | S |
| AR-0091 | Serratia marcescens | R | S | S | S | S |
| AR-0099 | Serratia marcescens | R | S | I | I | S |
| AR-0121 | Serratia marcescens | R | S | I | S | S |
| AR-0122 | Serratia marcescens | R | S | I | S | S |
| AR-0123 | Serratia marcescens | R | S | I | S | S |
| AR-0124 | Serratia marcescens | R | S | I | S | S |
| AR-0130 | Serratia marcescens | R | S | R | S | S |
| AR-0131 | Serratia marcescens | R | S | R | S | S |
| AR-0517 | Serratia marcescens | R | R | R | R | R |
| AR-0520 | Serratia marcescens | R | R | I | R | R |
| AR-0521 | Serratia marcescens | R | R | R | R | R |
| AR-0608 | Serratia marcescens | R | I | R | R | S |
| UF-166 | Serratia marcescens | R | R | R | S | R |
| UF-263 | Serratia marcescens | R | S | R | S | R |
| UF-269 | Serratia marcescens | R | S | R | S | R |

**Table 2: MICs and susceptibilities of *Serratia marcescens* strains.** Minimal inhibitory concentration (MIC) ug/mL, pattern of susceptibility resistant (R), intermediate (I), and susceptible (S).

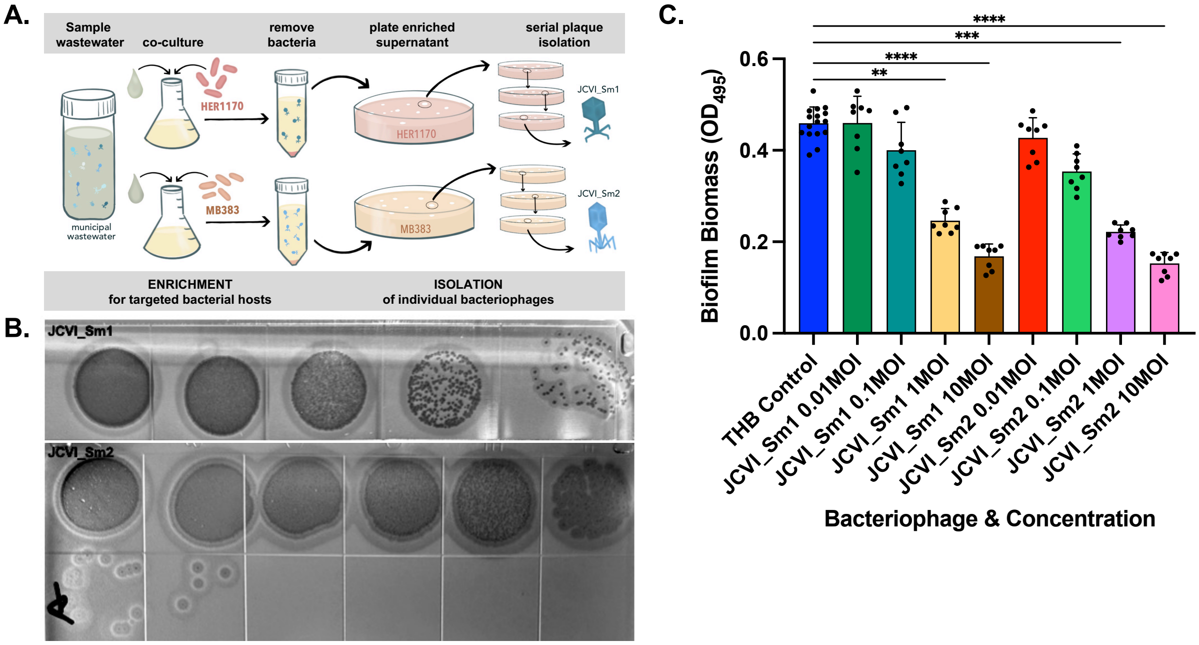

**Supplemental Figure 1**: **Isolation and Effect of bacteriophages in *S. marcescens* soft-agar cultures and mature biofilms**. **A**. Visual schematic of bacteriophage isolation from wastewater**. B.** Plaque assay for bacteriophages ϕJCVI_Sm1 and ϕJCVI_Sm2. **C** *Serratia marcescens* UT-383 was grown in biofilm form for 48 hours prior to ϕJCVI_Sm1 or ϕJCVI_Sm2 at MOIs 0.01, 0.1, 1 and 10. Statistical differences were determined by the Kruskal–Wallis test with Dunn's multiple-comparison post-test. * $p \leq 0.05$, ** $p \leq 0.01$, *** $p \leq 0.001$, **** $p \leq 0.0001$. Error bars denote standard deviation.

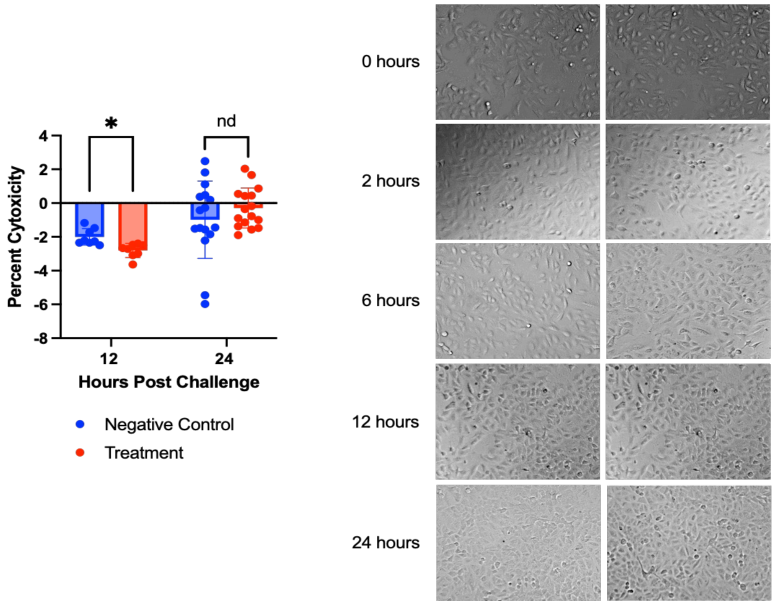

**Supplemental Figure 2**: **Sub-MIC antibiotics, AMPs and bacteriophage cocktail does not cause cytotoxicity in respiratory epithelial cells**. A549 cells were challenged with the Sub-MIC antibiotics, AMPs and bacteriophage cocktail and supernatant was collected at 12 and 24 hour to assess release of lactate dehydrogenase. Statistical differences were determined by Two-way ANOVA. * $p \leq 0.05$, ** $p \leq 0.01$, *** $p \leq 0.001$, **** $p \leq 0.0001$. Error bars denote standard deviation.
